# ProtFinder: An efficient machine learning framework for protein model selection on real data

**DOI:** 10.64898/2026.08.04.742760

**Authors:** Nguyen Huy Tinh, Yanghe Dong, Nhan Ly-Trong, Le Sy Vinh, Bui Quang Minh

## Abstract

Model selection is a fundamental step in phylogenetic analysis that determines the best-fit model of sequence evolution for a given multiple sequence alignment. Popular model selection methods, such as ModelFinder, rely on statistical information criteria, such as the Bayesian Information Criterion (BIC) or the Akaike Information Criterion (AIC). However, these approaches are computationally expensive and the use of information criteria has been the subject of ongoing discussion. Recently, machine learning has emerged as a promising approach for phylogenetic model selection in both nucleotide and protein sequence analyses. ModelDetector is currently the only machine learning-based method for amino acid substitution model selection. However, because ModelDetector was trained on simulated data, it does not perform well on real datasets. Another limitation is that it does not support different rate heterogeneity across sites (RHAS) models. To overcome these limitations, we introduce ProtFinder, an efficient machine learning framework for protein model selection that predicts amino acid substitution models, RHAS models, and amino acid frequency models. To enable ProtFinder to work with real datasets, we employed a transfer learning strategy consisting of three stages: (1) initial training on large-scale simulated data, (2) joint training on both simulated and real data, and (3) final fine-tuning using real data only. Experimental results show that ProtFinder outperformed ModelDetector in amino acid substitution model selection. ProtFinder achieved comparable accuracy to the maximum likelihood method ModelFinder for substitution model selection on medium and large MSAs. It performs slightly better than ModelFinder in RHAS model selection and substantially outperforms it in amino acid frequency model determination. Notably, ProtFinder is up to 1,400 times faster than ModelFinder in terms of inference time, making it particularly suitable for medium and large datasets.

## 1 Introduction

One of the most fundamental tasks in phylogenetic inference is to identify the best model to represent the evolutionary process underlying a given multiple sequence alignment (MSA) (Posada and Crandall, 1998; Abascal et al., 2005; Kalyaanamoorthy et al., 2017; Abadi et al., 2020; Burgstaller-Muehlbacher et al., 2023; Tinh and Vinh, 2025). Amino acid substitutions are typically modeled by a continuous-time, time-reversible, stationary Markov process represented by an instantaneous substitution rate matrix (Felsenstein, 2003). The matrix *Q* is the product of two components: an exchangeability rate matrix *R* and an amino acid frequency vector Π. The exchangeability rate matrix *R* is symmetric and comprises 189 free parameters that must be empirically estimated from large datasets with many protein MSAs. A number of general empirical amino acid substitution models have been estimated from diverse protein datasets and are widely used in phylogenetic analysis, including JTT (Jones et al., 1992), WAG (Whelan and Goldman, 2001), LG (Le and Gascuel, 2008), and Q.pfam (Minh et al., 2021). In addition, several clade-specific substitution models have been developed for particular taxonomic groups to better capture lineage-specific evolutionary patterns (Dang et al., 2010; Minh et al., 2021; Tinh and Vinh, 2024).

As the amino acid frequency vector Π contains only 19 free parameters, it can either be fixed to amino acid frequencies estimated from a large collection of MSAs (called the empirical amino acid frequency or -F model) or calculated directly from an input MSA under the study (called +F model).

To account for rate heterogeneity among sites (RHAS), where different positions in a sequence evolve at different rates (Yang, 1996), several rate models have been proposed and frequently used including the Invariable site model (+I) (Palumbi, 1989; Gu et al., 1995), the Gamma distribution model (+G) (Yang, 1994), and their combination (+I+G). Model selection methods usually rely on statistical information criteria, such as the Akaike Information Criterion (AIC) (Akaike, 1974) and the Bayesian Information Criterion (BIC) (Schwarz, 2007). For instance, the widely used model selection method ModelFinder (Kalyaanamoorthy et al., 2017) identifies the best-fit model for a given MSA by evaluating BIC scores of candidate models and selecting the one with the lowest BIC score. However, the information criterion-based approach requires repeated maximum-likelihood optimizations for each candidate model, making model selection very computationally expensive for large datasets. Moreover, several studies (Jhwueng et al., 2014; Seo and Thorne, 2018; Susko and Roger, 2020; Abadi et al., 2020; Crotty and Holland, 2022; Burgstaller-Muehlbacher et al., 2023) have questioned the applicability of information criteria in phylogenetic model selection because the assumptions underlying these criteria might be violated in phylogenetic analyses.

Machine learning approach has been used in different scientific fields, including phylogenetics. For nucleotide data, several machine learning-based methods have been proposed for nucleotide substitution model selection, including ModelTeller (Abadi et al., 2020) and ModelRevelator (Burgstaller-Muehlbacher et al., 2023). For protein data, ModelDetector (Tinh and Vinh, 2025) is currently the only machine learning-based method for amino acid substitution model selection. ModelDetector employs a neural network to identify the best-fit substitution model from a predefined set of empirical models. However, like the existing nucleotide-based methods, it was trained only on simulated datasets, and its applicability to real biological datasets requires validation. Another limitation of ModelDetector is its lack of support for RHAS model selection and amino acid frequency determination, both of which are routinely used in phylogenetic analyses.

To address these limitations, we introduce ProtFinder, an efficient machine learning framework for amino acid model selection. ProtFinder consists of three key components: (1) QFinder for selecting the best-fit exchangeability rate matrix; (2) FFinder for determining the amino acid frequency model (+F or -F model); and (3) RHASFinder for identifying the rate heterogeneity among sites model. To enable ProtFinder to analyze real biological data, we employed a transfer learning strategy. The neural networks were first pre-trained on a large simulated dataset, then trained jointly on both simulated and real datasets, and finally fine-tuned on real datasets to improve their performance and generalization. Experimental results on both simulated and real datasets demonstrate the effectiveness of ProtFinder in comparison with the maximum likelihood-based method ModelFinder and its superiority over the machine learning-based method ModelDetector.

## 2 Materials and Methods

ProtFinder comprises three classifiers: (1) QFinder, which selects the exchangeability rate matrix; (2) FFinder, which determines the amino acid frequency model; and (3) RHASFinder, which predicts the RHAS model. Figure 1 illustrates an overview of the ProtFinder workflow which includes four main stages: Data preprocessing, Network design, Network training, and Network evaluation. Each stage is described in the following sections.

**Figure 1.**
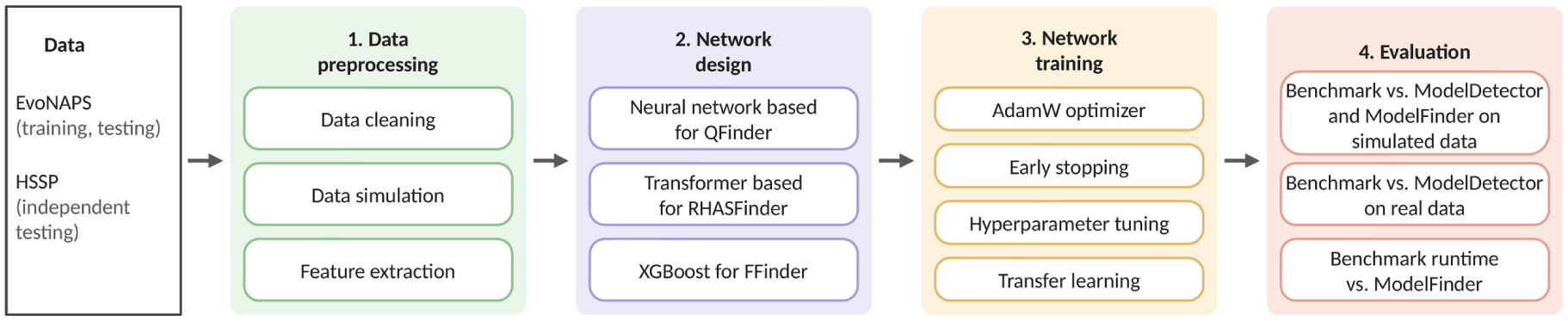
An overview of the ProtFinder workflow comprising four stages: Data preprocessing, Network design, Network training, and Network evaluation.

### 2.1 Data Pre-processing

#### Real databases

In this study, we used the EvoNAPS database (https://github.com/Cibiv/EvoNAPS), which combines MSAs from four published databases: PANDIT (Whelan et al., 2006), OrthoMaM (Allio et al., 2024), BenchmarkAlignments (https://github.com/roblanf/ BenchmarkAlignments), and TreeBASE (Sanderson et al., 1994). Specifically, EvoNAPS contains 21,800 protein alignments together with the corresponding best-fit models and phylogenetic trees inferred by IQ-TREE, spanning a wide range of species.

Among 40 amino acid substitution models available in IQ-TREE, seven models, comprising four general models (LG, WAG, JTT, and Q.pfam) and three clade-specific models (Q.bird, Q.mammal, and Q.plant), were identified as the best-fit models for 87% of all alignments in the EvoNAPS database. The remaining models were rarely selected, resulting in an insufficient number of empirical alignments for training and evaluating the classifiers. Therefore, we restricted our analysis to these seven models. To account for rate heterogeneity among sites, we considered four RHAS models: no rate heterogeneity (None), invariable sites (+I), discrete Gamma distribution (+G), and their combination (+I+G).

We filtered the EvoNAPS database to retain only MSAs whose best-fit substitution model was one of the seven selected substitution models and whose RHAS model was one of the four considered RHAS models (None, +I, +G, or +I+G). We obtained a dataset comprising 15,330 MSAs. Among these, 88.4% used the empirical amino acid frequencies specified by the best-fit substitution models, whereas the remaining 11.6% used the amino acid frequencies estimated directly from the input MSA (with +F in IQ-TREE). The proportions of alignments assigned to the None, +I, +G, and +I+G models were 1.68%, 1.73%, 33.47%, and 63.12%, respectively.

To evaluate the generalization performance of ProtFinder and ModelDetector, we tested both methods on an independent real HSSP dataset (Schneider et al., 1997) comprising 1,471 MSAs. The best-fit models inferred by ModelFinder were used as the reference labels for evaluation.

Figure 2 summarizes the characteristics of the two real datasets. In the EvoNAPS dataset, most MSAs contain fewer than 2,000 sites, with only 10 MSAs exceeding 10,000 sites. In the HSSP dataset, the number of taxa ranges from 10 to 285, while the alignment length ranges from 48 to 1,131 sites. On average, EvoNAPS contains more taxa per MSA than HSSP, with mean numbers of 84 and 59 taxa, respectively. This indicates that the HSSP testing dataset differs from the EvoNAPS training dataset, providing an independent evaluation of ProtFinder’s generalization ability.

**Figure 2.**
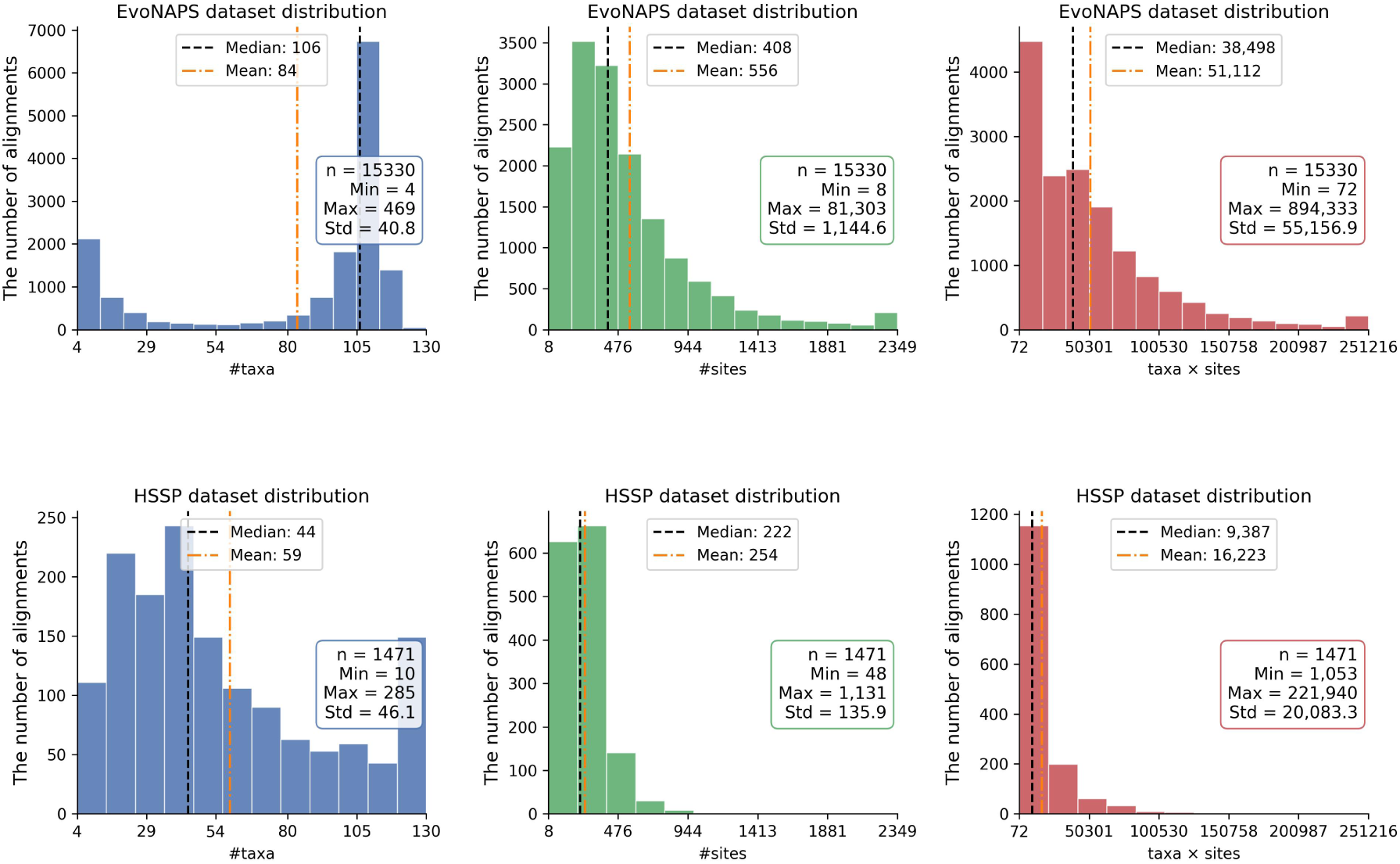
Distribution of the EvoNAPS and HSSP datasets by the number of taxa, alignment length, and dataset size (taxa × sites).

#### Training, Validation and Testing Datasets

We simulated a number of datasets to train our neural networks. Instead of sampling simulation parameters from predefined distributions, we estimated the empirical cumulative distribution function (ECDF) of each parameter from the EvoNAPS dataset and used inverse transform sampling to generate realistic parameter values. Specifically, for each parameter, a uniform random variable *u* ∼ U(0, 1) was sampled, and the corresponding value was obtained as

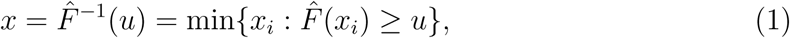

where *F̂* denotes the ECDF. This approach generates simulated data that reflect the empirical distributions of the real data.

We employed AliSim (Ly-Trong et al., 2023) to simulate protein MSAs using the simulation settings summarized in Table 1, including branch lengths, amino acid frequencies, the proportion of invariable sites, and the Gamma shape parameter *α*. This design resulted in 1,344 different parameter combinations (7 substitution models × 4 RHAS models × 2 amino acid frequency options × 6 tree sizes × 4 alignment lengths). All simulated MSAs were generated without gaps.

**Table 1:** Simulation settings.

| Settings | Options | Remarks |
| --- | --- | --- |
| Substitution model | LG, WAG, JTT, Q.plant, Q.bird, Q.mammal, Q.pfam |  |
| Amino acid frequency model | +F or -F. | - ‘+F’: amino acid frequencies are directly calculated from the input MSA;<br>- ‘-F’: using empirical amino acid frequencies specified by substitution models. |
| RHAS model | None, +I, +G, +I+G |  |
| Tree size | 8, 16, 32, 64, 128, 256 taxa | Phylogenetic trees were generated under the Yule-Harding model (Yule, 1925; Harding, 1971) with internal and external branch lengths randomly sampled from the empirical distributions estimated from the EvoNAPS dataset. |
| Alignment length | 100, 500, 1000, 3000 sites |  |

The training process of ProtFinder consists of three stages: (1) initial training on large-scale simulated data, (2) joint training on both simulated and real data, and (3) final fine-tuning using real data only. For the initial training stage, we simulated 200 MSAs for each parameter combination, yielding a total of 268,800 simulated MSAs. These alignments were randomly divided into training and validation sets with a ratio of 80% and 20%, respectively. For the joint training stage, we simulated 10 additional MSAs for each parameter combination, generating 13,440 MSAs, which were combined with the 15,330 real alignments in the EvoNAPS dataset. Finally, ProtFinder was fine-tuned using the 15,330 real alignments from the EvoNAPS dataset.

To evaluate ProtFinder on simulated data, we generated an independent test dataset comprising 53,760 simulated MSAs (40 MSAs for each parameter combination). We further evaluated ProtFinder and ModelFinder on the independent HSSP dataset, comprising 1,471 real MSAs, to assess their agreement on real data.

### 2.2 Model Selection using Maximum Likelihood Method

Given a multiple sequence alignment *D* and a set of amino acid substitution models Q, a set of RHAS models R, the maximum likelihood model selection method such as ModelFinder determines a tree *T*, a model *Q* ∈ Q and a model *R* ∈ R to maximize the likelihood value *L*(*Q, R, T* | *D*). It selects the best-fit model using the information criteria such as Bayesian Information Criterion (BIC) (Schwarz, 2007), which penalizes model complexity according to the number of free parameters *d*.

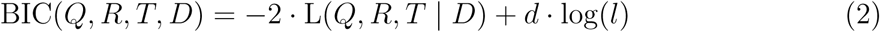

where *l* is the alignment length. A lower BIC score indicates a better-fit model.

Searching for the phylogenetic tree *T*, substitution model *Q*, and RHAS model *R* simultaneously is computationally expensive and only feasible for small alignments. The approximate method has been proposed in previous studies (Whelan and Goldman, 2001; Le et al., 2012; Dang et al., 2014; Minh et al., 2021) to overcome the computational burden. Specifically, ModelFinder first quickly approximates the maximum likelihood tree *T* ^Fast^ using the general substitution model LG and RHAS model +G. Given the fixed tree *T* ^Fast^, it then selects the best-fit model *Q*\* ∈ Q and *R*\* ∈ R that minimizes the BIC score:

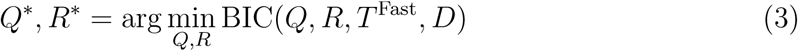

### 2.3 Feature Extraction

We extracted different sets of informative features from each alignment *A* to serve as input to three classifiers of ProtFinder.

For QFinder model, we randomly sampled 625 sequence pairs with replacement from each alignment *A*. This sampling size was selected based on benchmarking experiments using 225, 625, and 1,225 sequence pairs, where 625 achieved the best balance between prediction accuracy and computational efficiency. For each sequence pair, we extracted 440 normalized statistical features, including substitution frequencies among amino acids, amino acid invariability counts, and amino acid frequencies (see Table 2 for details). All features were normalized by the sequence length.

**Table 2:** Statistical features extracted for QFinder.

| Feature Name | Description | #Features |
| --- | --- | --- |
| Amino acid substitution frequencies | For each pair of amino acids $i$ and $j$ ($i \neq j$), counting the number of times that $i$ appears in sequence 1 and changes to $j$ in sequence 2. | 380<br>(i.e., $20 \times 19$) |
| Amino acid invariable site count | For each amino acid, counting the number of sites where the amino acid remains unchanged from sequence 1 to sequence 2. | 20 |
| Amino acid frequencies | For each sequence, counting the number of occurrences of each amino acid. | 40<br>(i.e., $2 \times 20$) |

The extracted features formed a 440 × 625 feature matrix, which was subsequently reshaped into a 440 × 25 × 25 tensor for input to the QFinder model. These features were specifically designed to capture the key characteristics underlying substitution model selection, including amino acid exchangeability patterns and equilibrium amino acid frequencies, the two fundamental components of amino acid substitution models.

For FFinder model, we extracted following features for each alignment *A*: the observed amino acid frequencies in *A*, the Kullback–Leibler divergence (KLD), the Jensen–Shannon divergence (JSD), minimum divergence values, and distances from divergences to their corresponding minimum values. Specifically, let *O_A_* = (*o*_1_*, . . ., o*_20_) denote the observed amino acid frequencies counted from MSA *A*. For each amino acid substitution model *Q*, let Π*_Q_* = (*π*_1_*, . . ., π*_20_) denote the empirical amino acid frequencies of *Q*. The Kullback–Leibler divergence between MSA *A* and model *Q*, based on amino acid frequency distributions, is computed as:

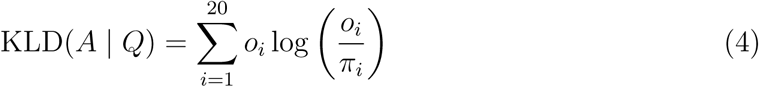

As KLD is asymmetric, we additionally computed the JensenShannon divergence (JSD), a symmetric and smoothed version of KLD, to measure the discrepancy between the amino acid frequency distributions. JSD is defined as:

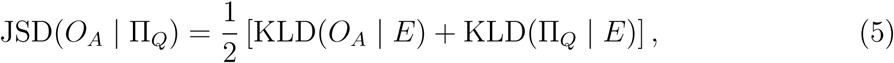

where

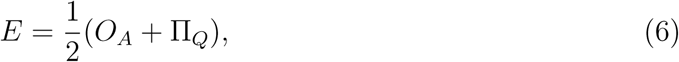

A low average JSD indicates that the amino acid frequencies in *A* are generally consistent with the empirical amino acid frequency distribution of the candidate substitution model *Q*. Therefore, JSD provides an informative feature for distinguishing amino acid frequency patterns.

In summary, FFinder uses a total of 50 features, including 20 observed amino acid frequencies, seven KLD values and seven JSD values (one for each candidate substitution model), the number of taxa, the number of sites, and 14 features representing the differences between each divergence value and its corresponding minimum divergence value. These features are used to determine whether amino acid frequencies should be estimated from the input MSA or taken from the substitution model.

For RHASFinder model, we extracted 23 statistical features for every site *i* in alignment *A* as below:

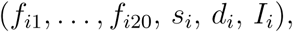

where

- *f_ij_* is the relative frequency of amino acid *j* (*j* = 1*, . . .,* 20) at site *i*,
- 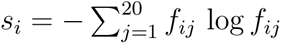 is the Shannon entropy,
- 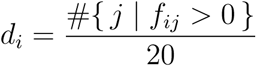 is the normalized diversity count,
- *I_i_* = 1 if *s_i_* = 0 and *I_i_* = 0 otherwise.

The site-specific Shannon entropy *s_i_* serves as a robust indicator of the substitution rate at site *i*; *d_i_* and *I_i_* summarize the amino acid variability at that site. Stacking the feature vectors from all sites shapes a *l* ×23 feature matrix where *l* is the alignment length. Ablation experiments indicated that removing any of the four feature groups {*f_ij_*}, *s_i_*, *d_i_*, or *I_i_* considerablly degraded the accuracy of the network, highlighting the complementary contributions of all feature groups.

To provide a more global alignment statistics for RHASFinder, we included 10 additional summary features: the proportion of invariable sites, the proportions of sites with *s_i_ <* 0.01 and *s_i_ <* 0.1, the 10^th^ and 25^th^ percentiles, the mean of ratio of gap sites, the variance of entropy, the skewness, the kurtosis, and the bimodality coefficient of the site entropy distribution.

To handle gaps in real alignments, we evaluated two strategies: replacing each gap with the most frequent amino acid at the corresponding site, or excluding gapped sites in the feature extraction. Benchmarking experiments showed that gap replacement achieved the highest accuracy for QFinder, whereas excluding gapped sites yielded superior performance for both RHASFinder and FFinder. Therefore, the optimal strategy was adopted for each classifier in all subsequent analyses.

### 2.4 Network Design and Implementation

We explored both neural networks and XGBoost to identify the most appropriate design for each of the three tasks (i.e., amino acid substitution model selection, amino acid frequency determination, and RHAS prediction). For neural networks, we evaluated multiple architectures, such as fully connected networks (Lecun et al., 2015), convolutional networks (LeCun et al., 1998), LSTM (Hochreiter and Schmidhuber, 1997), ResNet (Sarwinda et al., 2021), Transformers (Vaswani et al., 2017), and combinations of these architectures, incorporating modifications to individual layers, to determine the optimal network architecture for each task.

Figure 3 illustrates the architecture of QFinder. The network begins with an input layer that reshapes the input data from (625,440) to (440,25,25), followed by four repeated units, each consisting of a convolutional block and a squeeze-and-excitation (SE) block (Hu et al., 2019). Each convolutional block contains a convolutional layer followed by a batch normalization layer and a ReLU activation. The SE block receives an input with 32, 64, or 96 channels and applies global average pooling to compress each channel into a single scalar. This compressed representation is passed through two fully connected layers with ReLU and sigmoid activations to generate channel-wise weights, which are then multiplied element-wise with the original input. After the four convolutional and SE units, an average pooling layer reduces the spatial dimension. The resulting tensor is flattened and fed into a fully connected layer to produce the probabilities of the seven amino acid substitution models.

**Figure 3.**
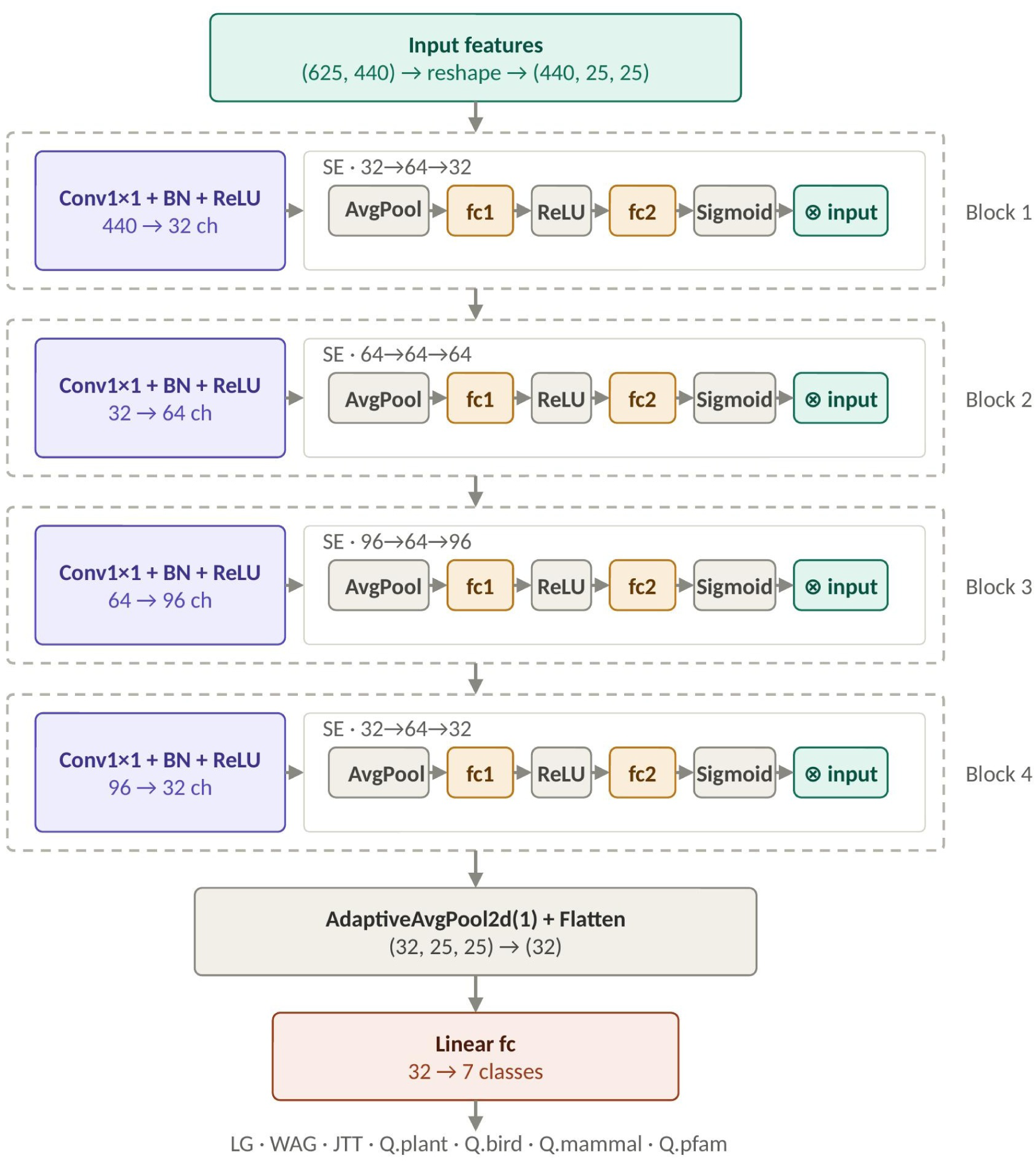
The network architecture of QFinder.

For FFinder, we used XGBoost with hyperparameters optimized by grid search to classify two labels: +F (estimating amino acid frequencies from the input alignment) and -F (using the amino acid frequencies specified by the substitution model).

Figure 4 outlines the network architecture of RHASFinder. First, the extracted site features are passed through a linear projection layer, after which the resulting feature sequence is processed by a Transformer encoder that stacks identical blocks of self-attention and position-wise feed-forward networks. The mean pooling is then applied across all sites to aggregate the token representations into a single embedding for the entire alignment. This embedding is concatenated with the global alignment statistics vector and passed through two fully connected layers with a ReLU activation to produce the logits for the four RHAS classes.

**Figure 4.**
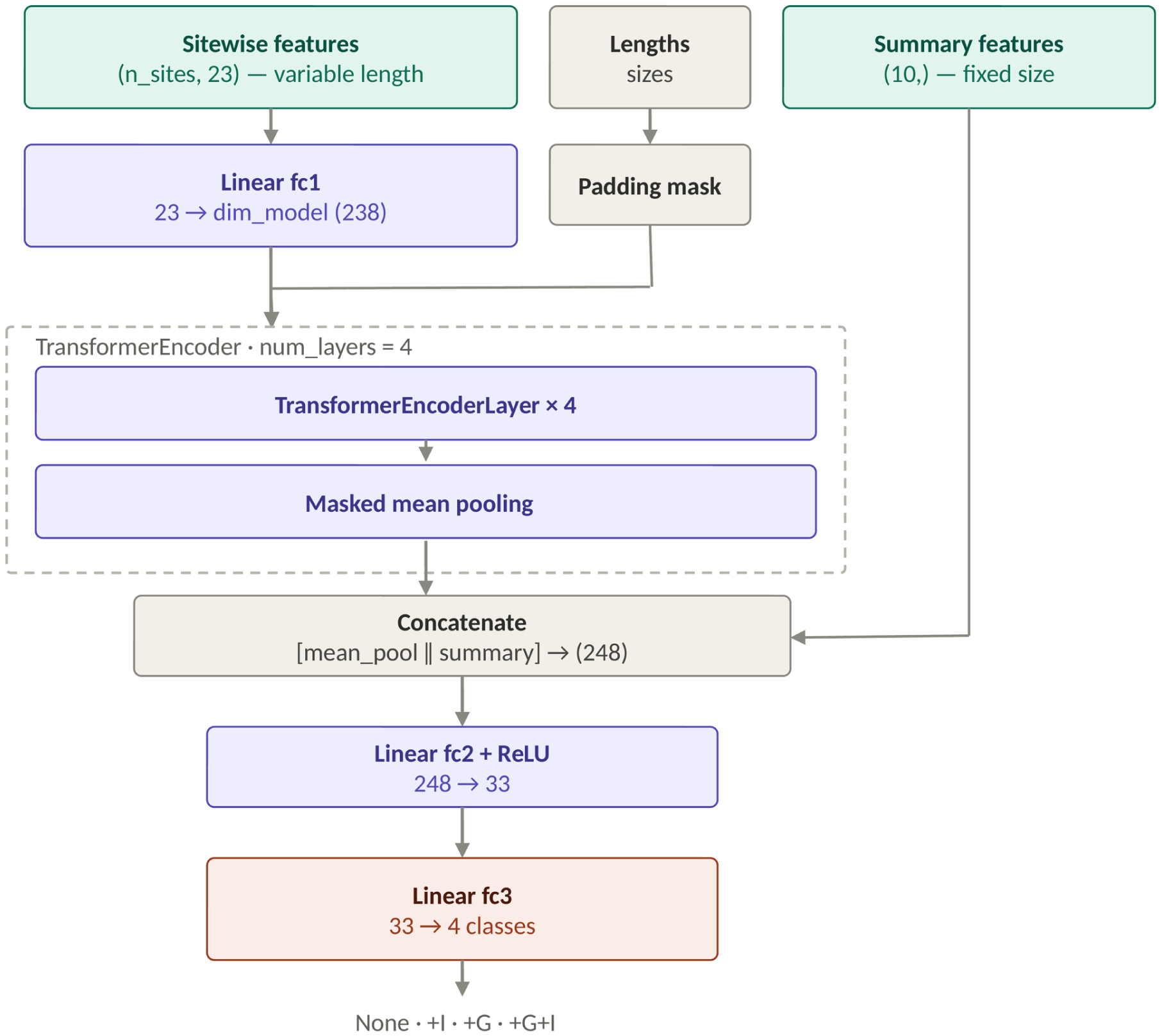
The network architecture of RHASFinder with main components.

Both QFinder and RHASFinder networks were trained with the AdamW optimizer (Loshchilov and Hutter, 2019). We also applied L2 regularization to mitigate overfitting. The networks were implemented in Python using PyTorch 2.8 with Automatic Mixed Precision (AMP) (Narang et al., 2018) employed to accelerate the training process and reduce memory usage. Hyperparameter tuning for QFinder and RHASFinder was carried out with Optuna (Akiba et al., 2019), an automated optimization framework that explored preset ranges of hyperparameters, including batch size, learning rate, and architecture specific variables.

### 2.5 Network Training

Since the amount of real data is limited, we employed a transfer learning strategy to train the classifiers. First, the classifiers were pre-trained on a large simulated dataset comprising 268,800 alignments. The pre-trained classifiers were then jointly trained on a combination of 13,440 simulated and 15,330 real alignments, and finally fine-tuned using the 15,330 real alignments only, with the initial learning rate reduced to one-tenth of its original value.

One major challenge with real data is class imbalance, which tends to bias predictions toward majority classes. We mitigated this issue by weighting the loss for each class with the reciprocal of its sample count. This increases the penalty for errors on rare classes and encourages the optimizer to treat all labels more uniformly. We also further evaluated two additional strategies: using weights as the reciprocal of the effective number of samples instead of raw counts, and replacing cross entropy with focal loss, but neither modification improved performance.

To prevent overfitting, we applied an early stopping mechanism that halted the training process if the validation accuracy did not improve by at least 0.2% for the last 10 consecutive epochs. During the initial training on simulated data, QFinder converged after 44 epochs, while RHASFinder converged after 34 epochs. In the joint training process on both simulated and real data, QFinder and RHASFinder converged after 47 and 41 epochs, respectively. Finally, fine-tuning on the real data required 35 epochs for QFinder and 31 epochs for RHASFinder.

Unlike the neural network-based models, FFinder employed XGBoost with a continued boosting mechanism, in which the booster obtained from one training stage was used to initialize the next stage. This enabled progressive adaptation to empirical data while retaining the knowledge learned in previous stages.

### 2.6 Evaluation

We evaluated the performance of ProtFinder on 53,760 simulated MSAs as well as 1,471 real MSAs from the independent HSSP database (Schneider et al., 1997). We compared its performance with the maximum likelihood method ModelFinder and our previous neural network ModelDetector (Tinh and Vinh, 2025).

ProtFinder supports seven substitution models and four RHAS models so we restricted ModelFinder to the same models by using commands ‘-mset LG,WAG,JTT,Q.bird,Q.mammal,Q.plant,Q.pfam’ for substitution models and ‘-mrate E,I,G,I+G’ for RHAS models. Note that ModelDetector was trained only for substitution model selection, therefore, we did not evaluate its performance for frequency or RHAS model selection.

For simulated datasets, the prediction accuracy of three methods was evaluated by comparing the inferred models with the true models used to generate the data. For real MSAs, where the true models are unknown, we benchmarked the predictions of ProtFinder and ModelDetector against those of ModelFinder.

## 3 Results

### 3.1 Performance Comparison on the Simulated Dataset

#### Overall Accuracy

Figure 5 compares the accuracies of ModelDetector, the three classifiers of ProtFinder (QFinder, FFinder, RHASFinder), and ModelFinder on simulated MSAs with different alignment lengths and numbers of taxa.

**Figure 5.**
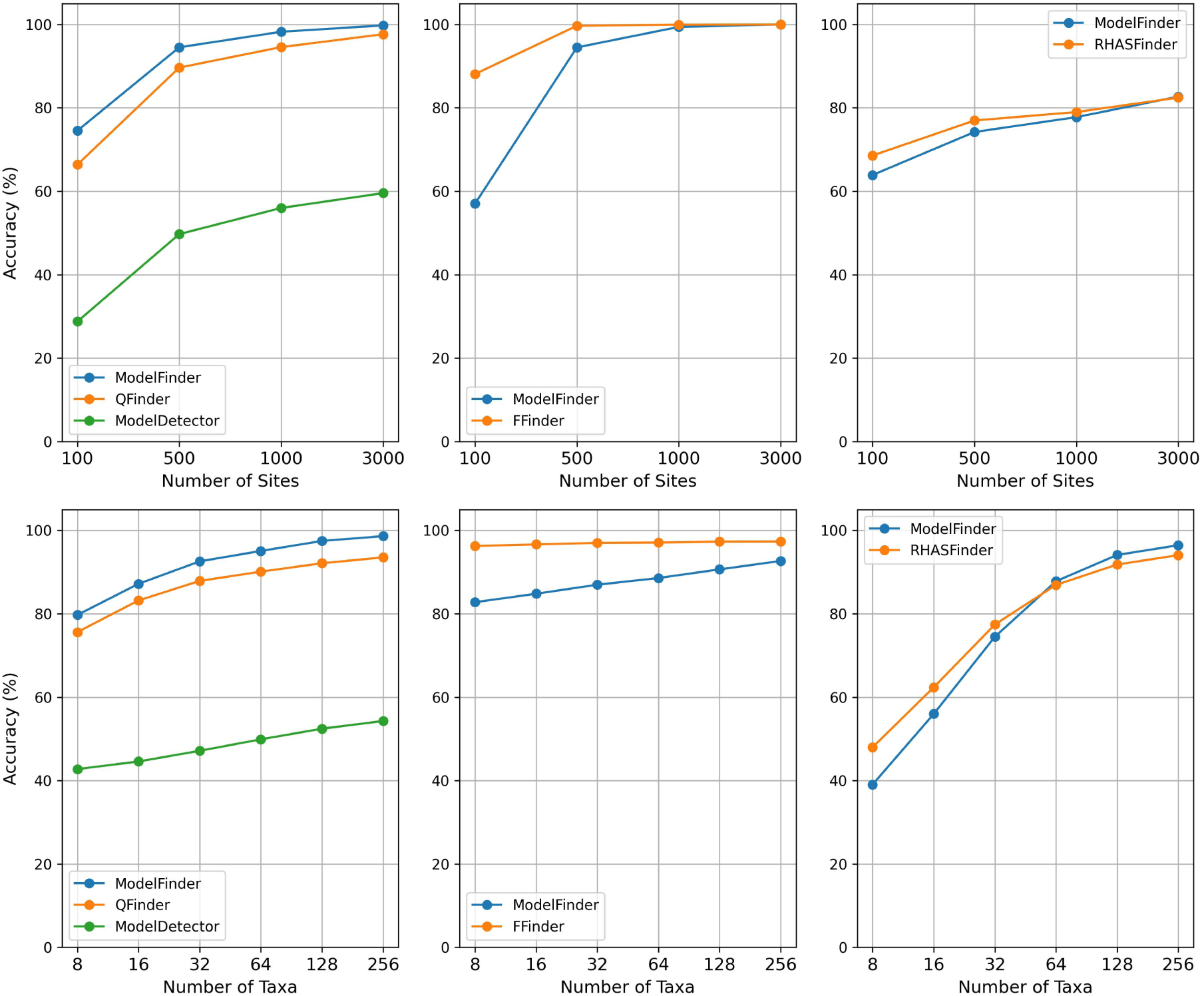
Average accuracy of ProtFinder, ModelDetector and ModelFinder on simulated testing data with different alignment lengths and numbers of taxa.

For the substitution model selection, on average, ModelFinder achieved the highest accuracy (91.75%), followed closely by QFinder (87.06%), whereas ModelDetector performed substantially worse (48.50%). Most misclassifications occurred in alignments with few taxa or short sequences, where limited phylogenetic signals make model discrimination more difficult for all methods. The accuracy of all three methods increased with alignment length and the number of taxa. For alignments containing 3,0000 sites, ModelFinder achieved an accuracy of 99.76%, while QFinder reached 97.65%. The much poorer performance of ModelDetector is likely due to its limited training configuration (i.e., it was trained only on simulated alignments under the I+G4 RHAS model).

ProtFinder consistently outperformed ModelFinder in both frequency and RHAS model prediction. On average, FFinder achieved an accuracy of 96.93%, compared with 87.73% for ModelFinder. Even for short alignments (100 sites), FFinder maintained high accuracy (88.10%), substantially outperforming ModelFinder (57.02%). As the alignment length increased, FFinder consistently achieved accuracies exceeding 99%, reaching 99.99% for alignments with 3,000 sites.

For RHAS inference, ProtFinder achieved a slightly higher average accuracy (76.75%) than ModelFinder (74.63%). The advantage became more pronounced as alignment size decreased. For example, on MSAs with 500 (or 100) sites, RHASFinder achieved an accuracy of 76.98% (or 68.58%), compared with 74.21% (or 63.90%) for ModelFinder. On MSAs with 16 taxa, RHASFinder correctly predicted 62.37% whereas ModelFinder obtained an accuracy of 55.98%.

#### Confusion Matrices of ProtFinder and ModelFinder

Figure 6 presents the confusion matrices of QFinder and ModelFinder for the substitution model selection. Both methods achieved high classification accuracy overall. QFinder showed lower accuracy than ModelFinder for the Q.pfam model, primarily due to its difficulty in distinguishing between the LG and Q.pfam models, both of which were estimated from the Pfam database and exhibit highly similar amino acid substitution patterns. Similarly, Q.bird and Q.mammal were occasionally confused because their model coefficients are highly correlated (i.e., Pearson correlation *r* = 0.98). Specifically, 11.39% of alignments simulated under Q.mammal were classified as Q.bird, whereas 11.38% of Q.bird alignments were classified as Q.mammal.

**Figure 6.**
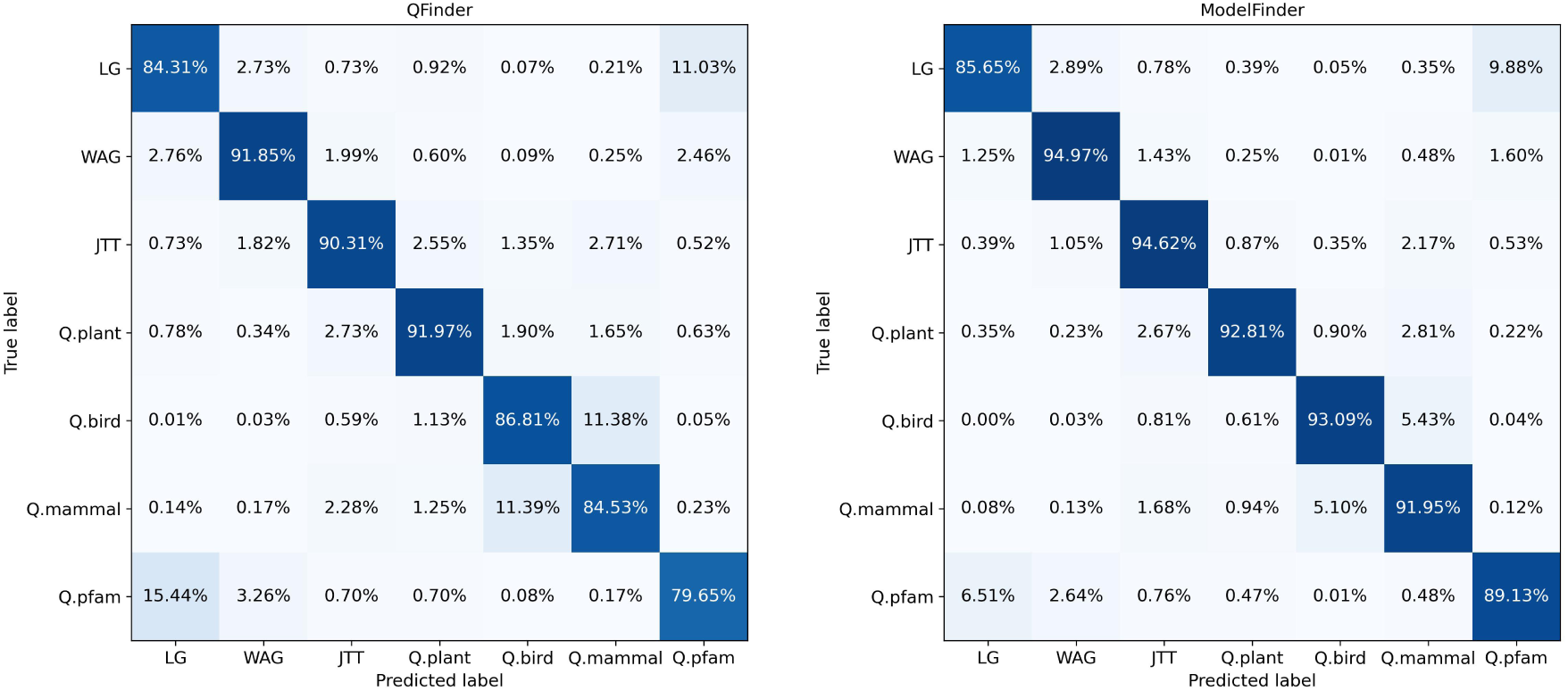
Confusion matrices of QFinder and ModelFinder on simulated testing data.

ModelFinder also exhibited similar confusion between pairs of highly correlated substitution models (i.e., Q.pfam and LG; Q.mammal and Q.bird), although these misclassifications were less pronounced than QFinder.

Figure 7 compares the confusion matrices of FFinder and ModelFinder for frequency model inference. Overall, FFinder shows more balanced performance for both +F and -F models. For the +F model, FFinder correctly classifies 97.24% of alignments, whereas ModelFinder misclassifies 24.55% of them. For the -F model, ModelFinder achieves 100% accuracy, while FFinder achieves 96.61%. These results suggest that ModelFinder is biased toward predicting the -F model, whereas FFinder provides a better balance between the two frequency models.

**Figure 7.**
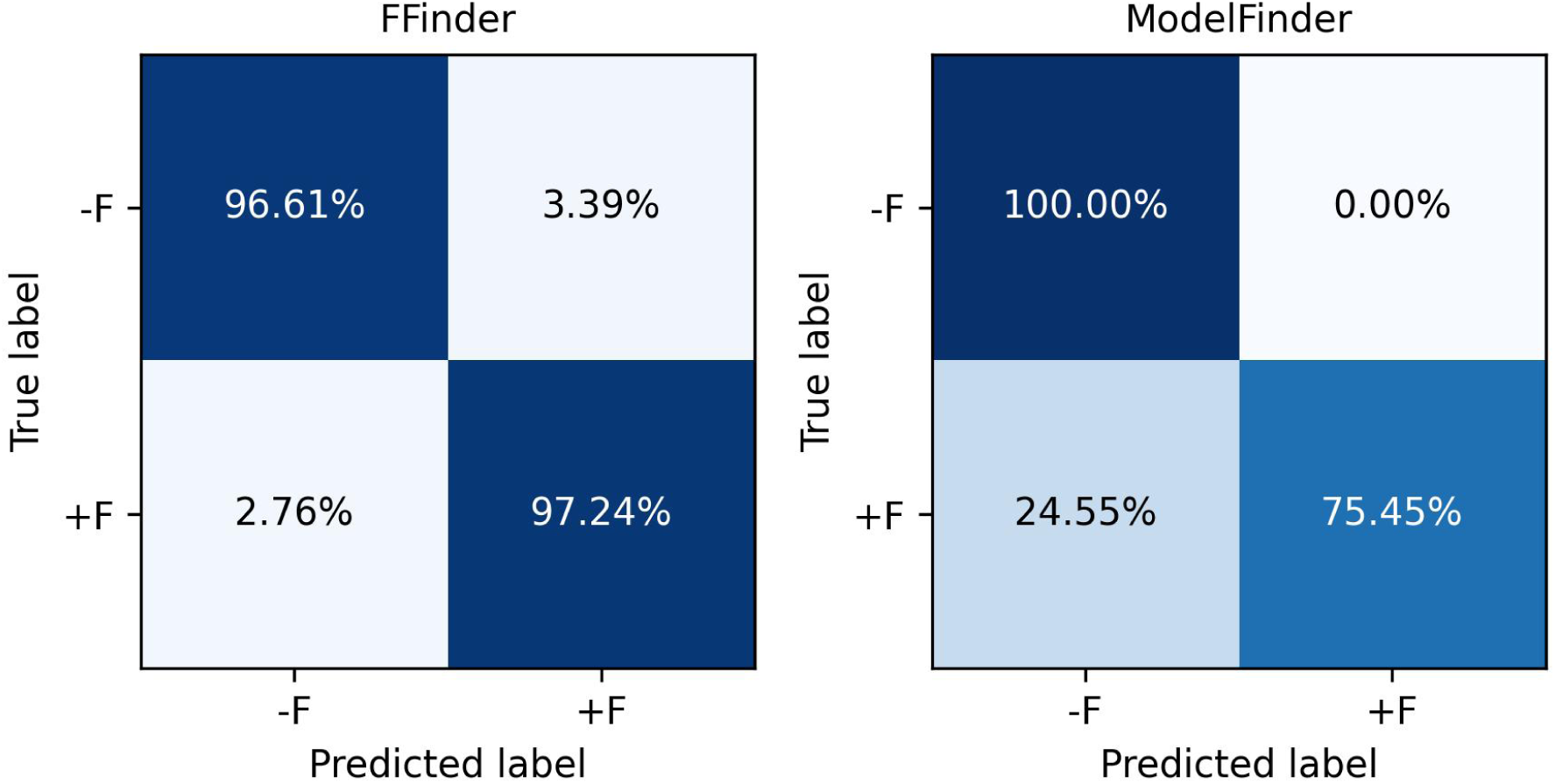
Confusion matrices of FFinder and ModelFinder on simulated testing data.

The confusion matrices in Figure 8 compare the performance of RHASFinder and ModelFinder for the RHAS model selection. Overall, RHASFinder demonstrates more balanced and accurate classification across all RHAS categories, whereas ModelFinder exhibits substantial confusion, particularly for models with more complex rate heterogeneity. For example, ModelFinder achievied an accuracy of only 45.04% for +I+G model, i.e., 40.6% of +I+G alignments are misclassified as +G, suggesting that ModelFinder often fails to detect the presence of invariant sites when both sources of rate heterogeneity coexist.

**Figure 8.**
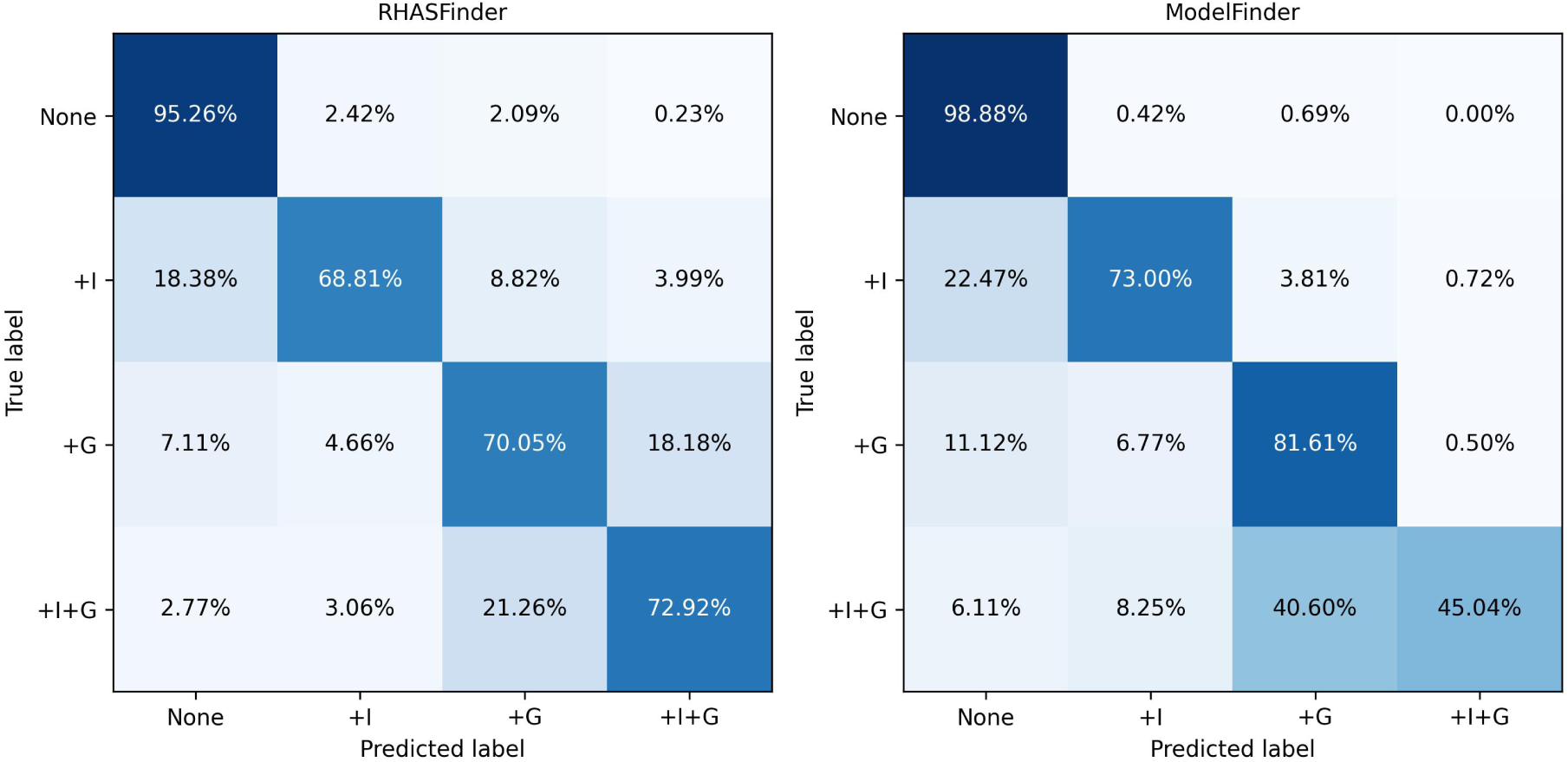
Confusion matrices of RHASFinder and ModelFinder on simulated testing data.

### 3.2 Performance Comparison on the Independent Real HSSP Dataset

As the true models are unknown for real MSAs, we used the best-fit models selected by ModelFinder as reference labels to evaluate the agreement of ProtFinder and ModelDetector with ModelFinder. We reported both top-1 agreement, where the predicted models of ProtFinder/ModelDetector matched the best-fit model selected by ModelFinder, and top-2 agreement, where the best-fit model selected by ModelFinder matched one of the two highest-probability models predicted by the machine learning methods. Because the dataset is imbalance, we report balance agreement and just considered groups that have at least 5% of total alignments. Note that since there are only two frequency model classes (+F and -F), top-2 agreement of FFinder is always 100%.

Figure 9 summarizes the top-1 and top-2 agreement of ModelDetector and ProtFinder (QFinder, FFinder, RHASFinder) with ModelFinder on real MSAs. FFinder achieved the top-1 agreement of 74.54%, followed by RHASFinder of 73.67% and QFinder of 67%. For substitution model selection, QFinder achieved a top-2 agreement of 90.96%, indicating high consistency with the best-fit models selected by ModelFinder. In comparison, ModelDetector achieved only 48.76% top-1 agreement and 70.77% top-2 agreement, likely because it was trained exclusively on simulated data under a fixed +I+G4 rate heterogeneity model. Similarly, RHASFinder achieved a top-2 agreement of 99.32%, indicating that one of its two highest-probability predictions was almost always the best RHAS model selected by ModelFinder. Overall, these results demonstrate that ProtFinder achieves high agreement with the likelihood-based ModelFinder on real MSAs.

**Figure 9.**
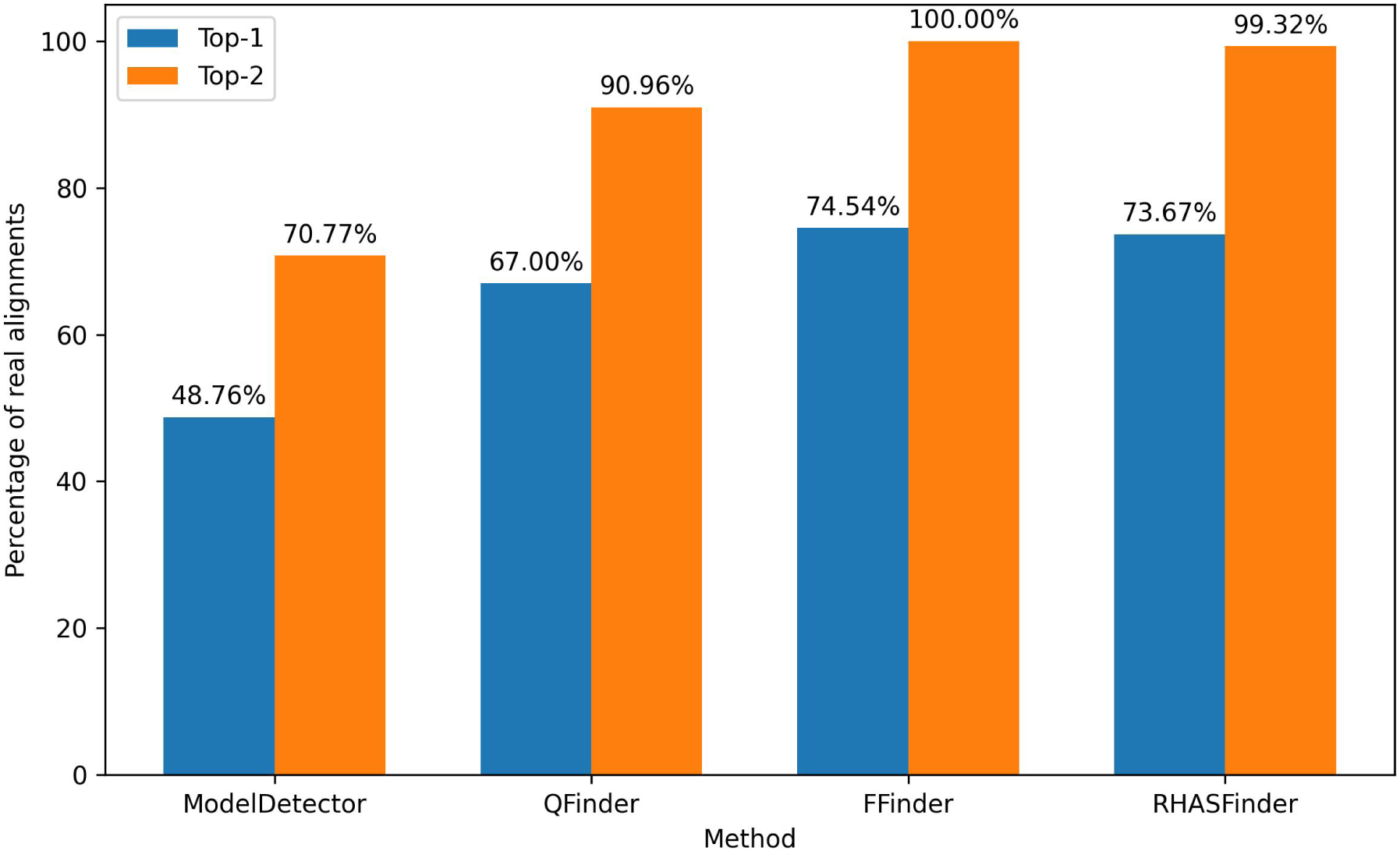
Agreement of ProtFinder and ModelDetector with ModelFinder on the independent real HSSP dataset. Top-1 denotes the percentage of alignments for which the model predicted by machine learning methods matches the best-fit model selected by ModelFinder. Top-2 denotes the percentage of alignments for which one of the two highest probability models predicted by machine learning methods matches the best-fit model selected by ModelFinder.

### 3.3 Time Analyses

The ProtFinder was trained on a machine with NVIDIA V100-PCIE-16GB GPU in less than 59 hours (10 hours for training QFinder, 48 hours for training RHASFinder and less than one hour for FFinder). Prediction time is the best advantage of machine learning methods over the maximum likelihood methods. All methods were executed on a single core of an Intel Xeon E5-2697 v4 2.3 GHz CPU.

The heatmap in Figure 10A shows the runtime ratio (ModelFinder/ProtFinder) in log-scaled, highlighting substantial speedups achieved by ProtFinder. The runtime of ProtFinder is defined as the total runtime of QFinder, FFinder, and RHASFinder, including feature extraction and model prediction. For simulated datasets with up to 256 taxa and 100 sites, ProtFinder is approximately 1, 400× faster than ModelFinder.

**Figure 10.**
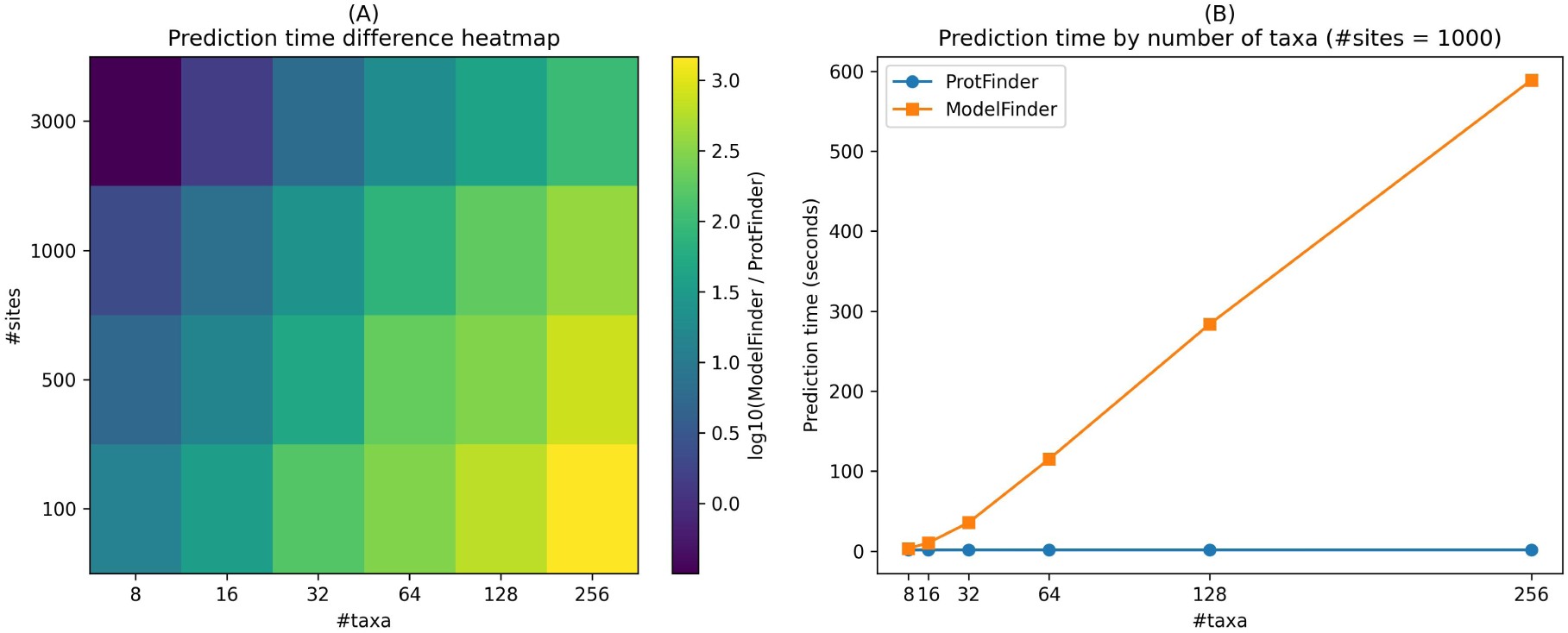
Runtime comparison between ProtFinder and ModelFinder. (A) Heatmap showing the log-scaled runtime ratio (ModelFinder/ProtFinder). (B) Runtime of ProtFinder and ModelFinder across varying numbers of taxa with a fixed sequence length of 1,000 sites.

While the runtime of ModelFinder increased rapidly with the number of taxa or alignment sites, the runtime of ProtFinder remained nearly constant (Figure 10B). For alignments with 256 taxa and 1,000 sites, ModelFinder required about 10 minutes to select the best-fit substitution and RHAS models, whereas ProtFinder completed the same task in only 1.5 seconds.

## 4 Conclusion and Discussion

Model selection is a fundamental step in phylogenetic inference. Traditional likelihood-based methods, such as ModelFinder, provide accurate model selection but are computationally expensive, particularly for large protein alignments. Machine learning methods, such as ModelDetector, substantially reduce prediction time but are trained exclusively on simulated data, limiting their ability to analyze real data.

In this study, we introduced ProtFinder, a machine learning framework trained on both simulated and real alignments for selecting amino acid substitution, frequency, and RHAS models. ProtFinder effectively combines different machine learning techniques, ranging from neural network (QFinder), transformer (RHASFinder) to XGBoost (FFinder). Across extensive benchmarking, ProtFinder consistently achieved high prediction accuracy, with accuracy improving as the numbers of taxa and alignment sites increased. For example, QFinder achieved an accuracy of substitution model selection greater than 95.27% for alignments containing at least 32 taxa and 1000 sites. These results suggest that ProtFinder is particularly well suited for medium to large protein alignments.

ProtFinder also offers substantial computational advantages over the maximum likelihood-based methods. Unlike ModelFinder, which evaluates candidate models sequentially, ProtFinder can process multiple alignments simultaneously through batch inference on modern hardware, reducing runtime by several orders of magnitude.

RHASFinder is based on a Transformer architecture, which requires substantial computational resources. Therefore, GPU acceleration is recommended to achieve efficient inference. Prediction time increases with alignment length because the Transformer’s computational cost scales with the number of sites. In our experiments, a NVIDIA Tesla V100-PCIE-16GB was able to process alignments of up to 15,000 sites without running out of memory, making it suitable for large-scale phylogenetic analyses.

The machine learning-based approach is less effective than the maximum-likelihood method at distinguishing between highly correlated substitution models because these models exhibit very similar amino acid substitution patterns. ProtFinder provides confidence scores that enable a practical two-stage model selection strategy. ProtFinder can first identify a small set of highly probable candidate models (e.g., the smallest set accounting for 95% of the cumulative prediction confidence), after which ModelFinder evaluates only these candidate models instead of a large number of available models. This strategy substantially reduces the computational cost of maximum-likelihood model selection while preserving its accuracy. We therefore plan to integrate ProtFinder into IQ-TREE as a fast pre-screening module to accelerate the model selection process.

## Acknowledgements

The authors thank Robert Lanfear, Thomas Wong, Piyumal Demotte, and Hashara Kumarasinghe for their valuable comments and insightful discussions. AI-assisted writing tools were used to improve spelling, grammar, and language during the preparation of this manuscript.

## Data Availability

All data, source codes and scripts used in the development of ProtFinder, including those for data generation, training, testing, and fine tuning QFinder, RHASFinder, and FFinder, are publicly available at https://github.com/tinhnh2/ProtFinder.

